# Non-coding sequence variation reveals fragility within interleukin 2 feedback circuitry and shapes autoimmune disease risk

**DOI:** 10.1101/2023.06.17.545426

**Authors:** Dimitre R. Simeonov, Kyemyung Park, Jessica T. Cortez, Arabella Young, Zhongmei Li, Vinh Nguyen, Jennifer Umhoefer, Alyssa C. Indart, Jonathan M. Woo, Mark S. Anderson, John S. Tsang, Ronald N. Germain, Harikesh S. Wong, Alexander Marson

**Author notes:** Equal authorship (Dimitre R. Simeonov, Kyemyung Park). Co-senior authorship and co-correspondence, Correspondence.

## Abstract

Genetic variants associated with human autoimmune diseases commonly map to non-coding control regions, particularly enhancers that function selectively in immune cells and fine-tune gene expression within a relatively narrow range of values. How such modest, cell-type-selective changes can meaningfully shape organismal disease risk remains unclear. To explore this issue, we experimentally manipulated species-conserved enhancers within the disease-associated *IL2RA* locus and studied accompanying changes in the progression of autoimmunity. Perturbing distinct enhancers with restricted activity in conventional T cells (Tconvs) or regulatory T cells (Tregs)—two functionally antagonistic T cell subsets—caused only modest, cell-type-selective decreases in *IL2ra* expression parameters. However, these same perturbations had striking and opposing effects *in vivo*, completely preventing or severely accelerating disease in a murine model of type 1 diabetes. Quantitative tissue imaging and computational modelling revealed that each enhancer manipulation impinged on distinct IL-2-dependent feedback circuits. These imbalances altered the intracellular signaling and intercellular communication dynamics of activated Tregs and Tconvs, producing opposing spatial domains that amplified or constrained ongoing autoimmune responses. These findings demonstrate how subtle changes in gene regulation stemming from non-coding variation can propagate across biological scales due to non-linearities in intra- and intercellular feedback circuitry, dramatically shaping disease risk at the organismal level.

---

The adaptive immune system must protect the host from threats, such as pathogens and tumours, while limiting autoimmunity. Achieving this objective requires the use of control circuits, particularly feedback regulation, which can dynamically steer the system’s output (i.e., the magnitude of an immune response) towards a desired target value over time and space. This concept is well exemplified by αβ T cells, which employ a multitude of such circuits spanning both intracellular and intercellular scales to rapidly amplify or dampen their inflammatory capacity(*1–6*). Consequently, T cells can convert relatively small changes in input signals into much larger changes in output responses— a non-linear behaviour enabling effective scaling in response to replicating pathogens or appropriate blunting of autoimmune responses.

The inherent nonlinearities within these control circuits suggest that minor imbalances in signal processing could lead to inappropriate scaling of self-reactive Tconv responses, which are initiated in LNs on an ongoing basis but generally constrained by Tregs in healthy hosts(*3, 7*). In support of this concept, both theoretical and experimental studies exploring the parameter space of different T cell control circuits have demonstrated robustness over a wide range of values for some parameters, yet substantial fragility outside of a narrow range for others (*2, 3, 8–10*). Even small changes in fragile parameters can have considerable effects on the magnitude of self-reactive Tconv responses, resulting in tissue damage(*3, 11, 12*). This paradigm has important implications for understanding the genetic basis of autoimmunity. Indeed, most common human autoimmune disease-associated variants map to non-coding enhancers and appear to exert relatively modest quantitative or kinetic effects on target gene expression, in contrast to complete loss- or gain-of-function mutations(*13, 14*). Moreover, enhancers harboring these non-coding variants can exhibit selective activity in specific immune cell types, especially T cell subsets(*13, 15–22*). We thus hypothesized that seemingly subtle enhancer variants impinging on fragile T cell circuit parameters could have much larger effects on organismal autoimmune disease risk.

To examine this concept, we focused on the *IL2RA* locus, which harbors numerous genetic variants associated with different human autoimmune disorders. *IL2RA* encodes the high-affinity receptor (IL-2RA) for IL-2, a cytokine that regulates the functionality and fitness of activated Tconvs and Tregs. These two T cell subsets exhibit an opposing, reciprocal relationship: activated Tconvs produce IL-2 and require it to enhance their proliferation, survival, and effector state, whereas Tregs cannot produce IL-2 yet require it to maintain their numbers and boost their immunosuppressive functions(*3, 23–27*). Thus, the same cytokine can counterintuitively amplify and dampen the immune response. The outcome of the response strongly depends on the extent of IL-2RA expression quantity by individual Tconvs or Tregs as this receptor increases the binding affinity for IL-2 by up to three orders of magnitude and further influences distinct IL-2 feedback circuits operating within and between both T cell subsets(*3, 5, 28*). Non-linearities in these circuits may amplify and propagate small changes in IL-2 production or sensing, significantly favoring activated Tconvs or Tregs. These considerations suggest that 1) the control circuits governing self-reactive Tconv responses may exhibit fragility with respect to subtle variations in *IL2RA* expression parameters, and 2) *IL2RA* enhancer variants with selective activity in Tregs or Tconvs may have significant yet opposing effects on the progression of autoimmune disease.

We first sought to identify distinct *IL2RA* enhancers with selective activity in primary human Tconvs or Tregs. Our previous CRISPRa screen revealed a species-conserved *IL2RA* enhancer, termed the CRISPRa responsive element (CaRE) 4 enhancer, that operates predominantly in Tconvs (Fig. 1a). This region appears to control the kinetics of *IL2RA* induction following activation of Tconvs through their T cell receptor (TCR)(*29*). To further examine this conclusion, we assessed histone 3 lysine 27 acetylation (H3K27Ac)—a cardinal feature of enhancer activity—by analyzing publicly available ChIP-seq data(*30*). This approach revealed a significant increase in H3K27Ac in the CaRE4 enhancer following activation of Tconvs, but not Tregs (Fig. 1b). Using the same ChIP-seq data set, we searched for candidate *IL2RA* enhancer elements with preferential activity in Tregs. We identified a distinct species-conserved sequence ∼200bp upstream of the *IL2RA* promoter that exhibited high baseline levels of H3K27Ac in Tregs, but not Tconvs (Fig. 1b). This region falls within a distinct CaRE affecting IL2RA protein expression in our previous CRISPRa screen, and therefore, we refer to it as the CaRE3 enhancer (Fig. 1a, b)(*29*). Notably, this enhancer exhibited selective binding to FOXP3 and STAT5, two transcription factors that directly promote Treg *IL2RA* expression(*27, 31, 32*). Unlike the CaRE4 enhancer, the CaRE3 enhancer did not display increased H3K27Ac following TCR-mediated activation (Fig. 1b). These results strongly suggest that the CaRE4 and CaRE3 enhancers selectively operate in activated Tconvs and resting Tregs, respectively.

**Figure 1.**
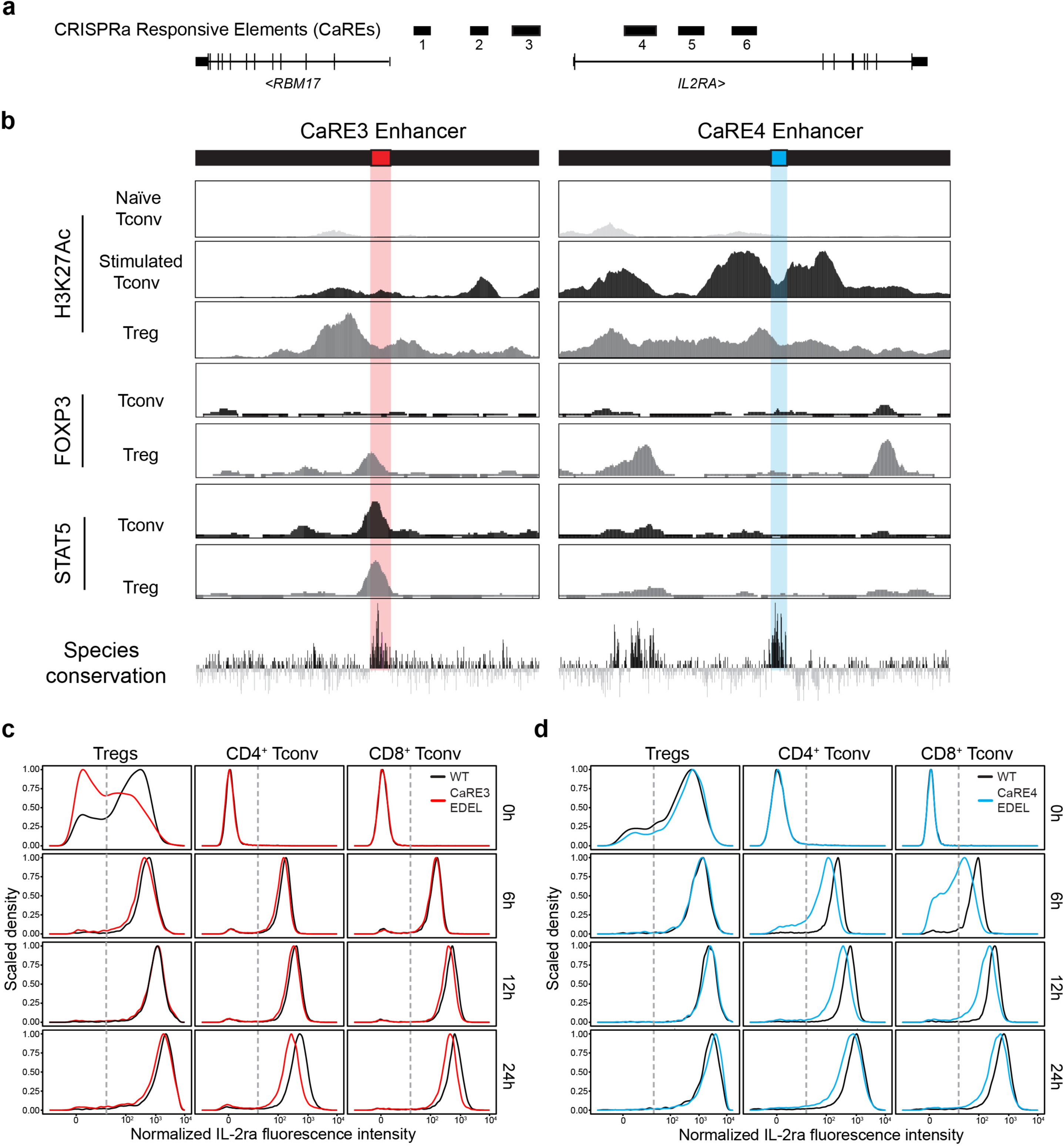
The CaRE3 and CaRE4 enhancers control distinct *IL2RA* expression parameters in Tregs and activated Tconvs. **a,** Schematic of the *IL2RA* locus and previously identified CRISPRa responsive elements (CaREs), including CaRE3 (red) and CaRE4 (blue)(*29*) **b,** ChIP-seq data for H3K27Ac and indicated transcription factors at the CaRE3 (hg19 chr10:6109198-6113209) and CaRE4 (hg19 chr10:6092205-6096760) regions in primary human T cell subsets. H3K27Ac data extracted from the Epigenome Roadmap Project(*30, 65*). H3K27Ac signal y-axis range 2-120 for CaRE3 and 2-300 for CaRE4. STAT5 and FOXP3 ChIP signal y-axis range 1-40 for CaRE3 and CaRE4. **c & d,** Representative histograms illustrating IL-2ra surface expression over time on stimulated Tconvs or Tregs from WT NOD or EDEL^+/+^ NOD animals. Cells were stimulated with high concentrations of anti-CD3/anti-CD28 *in vitro*.

Since both sequences are conserved across species, we proceeded to determine their influence on *IL2ra* expression parameters in primary murine T cells (Fig. 1c,d). For this purpose, we isolated splenocytes or naïve Tconvs from mice engineered with a germline CaRE4 or CaRE3 enhancer deletion (EDEL) and studied changes in T cell IL-2ra protein levels. The CaRE4-EDEL mice have been described previously whereas the CaRE3-EDEL mice were generated for this study, both in the C57BL/6 and non-obese diabetic (NOD) genetic backgrounds(*29*) (Supplemental Fig. 1 & 2). Importantly, deleting the CaRE3 region did not strongly alter the progression of T cell development *in vivo* as the abundance of T cell precursor populations (i.e., thymocytes) remained normal within the thymus compared to controls (Supplemental Fig. 1a, b & Supplemental Fig. 2b, c). We did, however, observe a reduction in IL-2ra protein expression on double-negative (DN) thymocytes—the earliest T cell precursor—along with changes in the abundance of progenitor Tregs (Supplemental Fig. 1c-e & Supplemental Fig. 2d-f). Despite these latter changes, the frequency of mature Tregs within peripheral blood and secondary lymphoid organs (SLOs) remained comparable between wild-type (WT) and CaRE3-EDEL mice (Supplemental Fig. 1f, g & Supplemental Fig. 2g, h).

**Figure 2.**
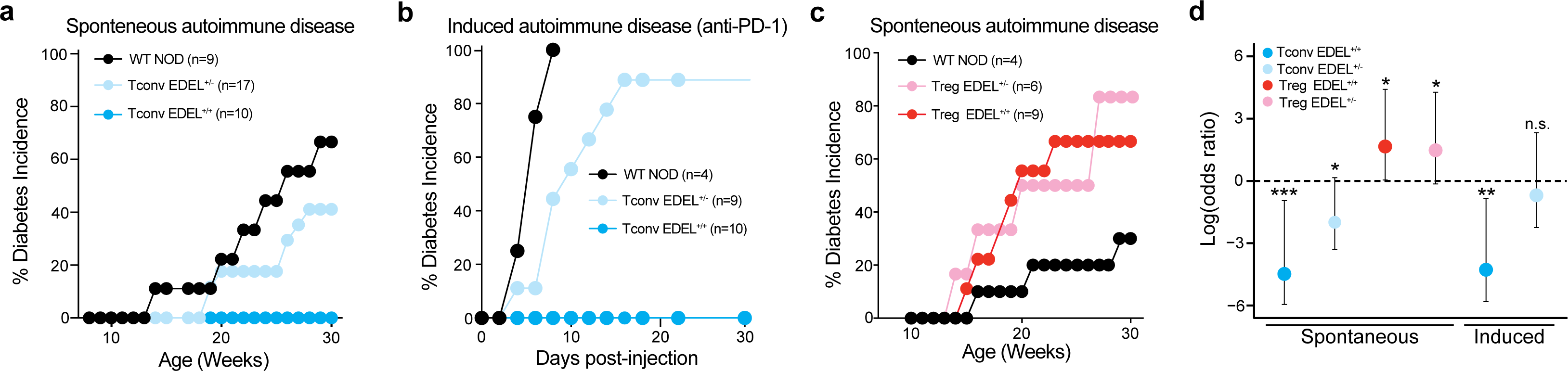
Small changes in Tconv or Treg *IL2ra* expression parameters due to CaRE3 or CaRE4 enhancer perturbations cause pronounced effects on autoimmune disease risk. **a**, Incidence of spontaneous autoimmune diabetes in female WT NOD or Tconv-EDEL NOD animals. **b,** Incidence of autoimmune diabetes following anti-PD1 treatment in female WT NOD or Tconv-EDEL NOD animals. **c,** Incidence of spontaneous autoimmune diabetes in female WT NOD or Treg-EDEL NOD animals. **d,** Natural logarithm of the odds ratios for developing spontaneous or induced autoimmune diabetes based on indicated genotypes compared to WT NOD littermate controls (dashed black line). Error bars represent 95% Wald confidence intervals computed using unconditional maximum likelihood estimation with a small sample size adjustment. Calculations were performed using the Haldane-Anscombe correction to avoid zero values in the 2x2 contingency table. Two-sided p-values were calculated using a chi-squared test.

We initially assessed IL-2ra protein expression in naive Tconvs, which must synthesize the receptor *de novo* following TCR-mediated activation, or Tregs, which express high levels of surface IL-2ra at steady-state, from wild-type (WT) NOD or EDEL NOD animals. Upon stimulation with high concentrations of anti-CD3/CD28 antibodies— a strong input signal known to result in significant amounts of IL-2 in the culture medium— activated Tconvs from WT and CaRE3-EDEL animals upregulated IL-2ra at comparable rates over time(*33*) (Supplemental Fig. 3a-c). By contrast, Tconvs from CaRE4-EDEL animals displayed a modest kinetic delay in IL-2ra expression levels, most strongly at early time points post-activation, consistent with our previous findings (Fig. 1c, d & Supplemental Fig. 3a, d, e).

**Figure 3.**
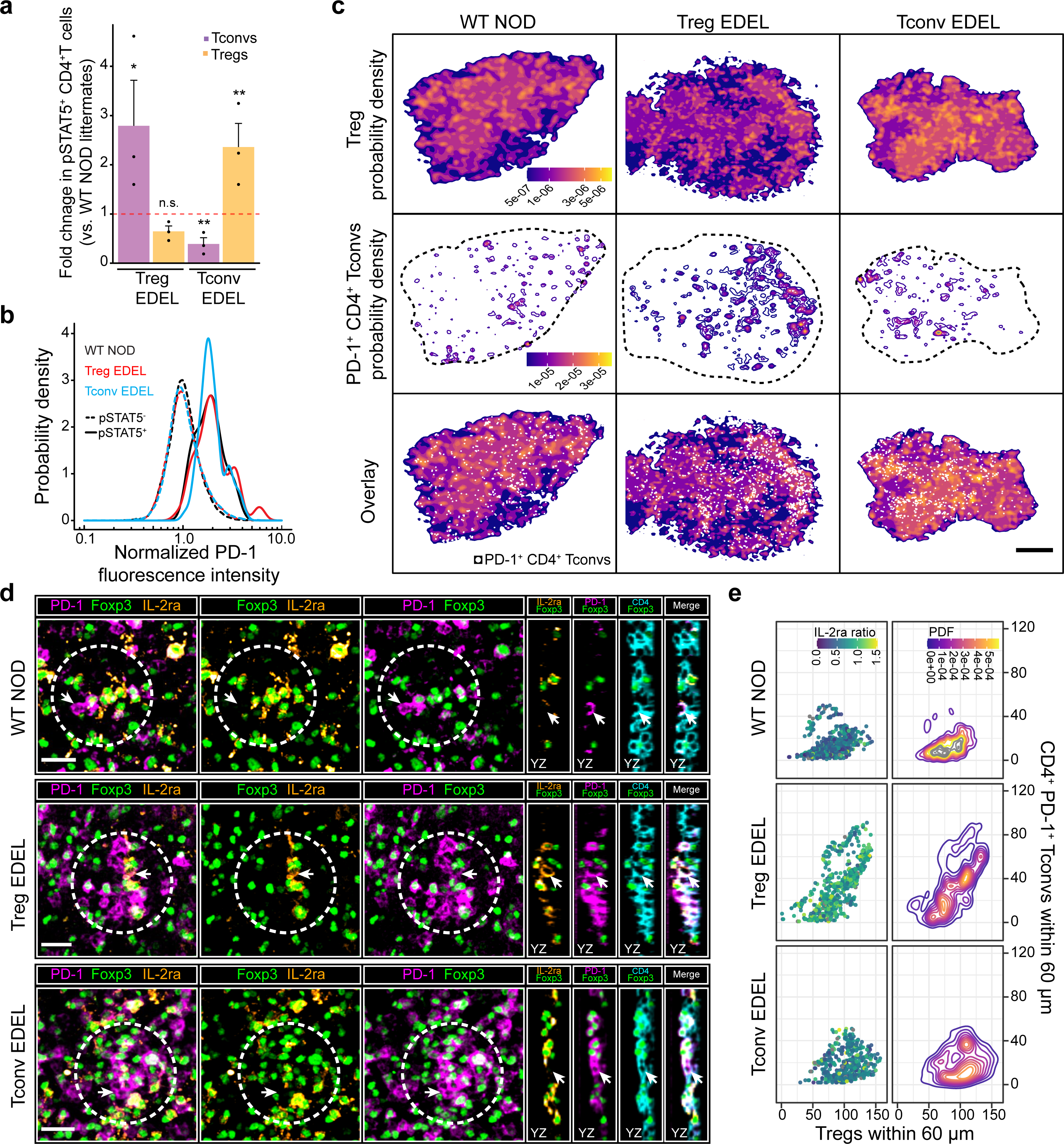
Persistent imbalances in Tconv or Treg IL-2 sensing result in opposing spatial domains that amplify or constrain ongoing autoimmune responses *in situ*. **a**, Fold-changes in pSTAT5^+^ CD4^+^ Tconvs or pSTAT5^+^ Tregs within the panLN paracortex of 6-week-old WT or EDEL^+/+^ NOD female mice. Individual dots represent individual mice. Data are presented as mean ± SEM from two independent experiments. * p-value = 0.05, ** p-value < 0.001 b, *In situ* quantification of PD1 expression on pSTAT5^-^ or pSTAT5^+^ CD4^+^ Tconvs. Analysis was spatially restricted to the panLN paracortex of WT or EDEL^+/+^ NOD mice. **c,** Spatial density functions of paracortical Tregs (top row) or PD-1^+^ CD4^+^ Tconvs (middle row) in the panLNs of WT or EDEL^+/+^ NOD mice. Bottom row, white dots represent the positions of individual PD1^+^ CD4^+^ Tconvs overlaid on top of paracortical Treg densities. Scale bar = 200 μm. **d,** Representative confocal micrographs illustrating changes in the local densities (dashed white circles) of Tregs and CD4^+^ PD1^+^ Tconvs as well as changes in their local IL-2ra expression ratios. Images acquired in the panLNs of WT or EDEL^+/+^ NOD mice. YZ optical slices highlight CD4, PD1, and IL-2ra expression on the cell of interest (white arrow). Scale bars = 25 μm. e, *In situ* multiparameter analysis of the number of Tregs (x-axis) and CD4^+^ PD1^+^ Tconvs (y-axis) surrounding individual CD4^+^ PD1^+^ Tconvs (individual dots). The IL-2ra ratio was calculated by dividing the median IL-2ra fluorescence intensity of each PD1^+^CD4^+^ Tconv by the median IL-2ra fluorescence intensity of the surrounding Tregs within 60 μm.

Tregs from WT and CaRE4-EDEL mice expressed comparatively high surface levels of IL-2ra at baseline and upregulated the receptor similarly following stimulation (Fig. 1d). This stimulation-responsive upregulation likely stems from 1) strong triggering of the Treg TCR and 2) Tregs responding to IL-2 released from activated Tconvs in the splenocyte culture—two signals that are well-known to enhance IL2ra expression(*5*). While Tregs from CaRE3-EDEL mice also expressed IL-2ra in the absence of stimulation, they exhibited reduced surface levels compared to Tregs from WT and CaRE4-EDEL mice (Fig. 1c). Specifically, in CaRE3-EDEL mice, we observed a 2-3-fold decrease in the fraction of Tregs with high IL-2ra, culminating in a ∼1.5-1.8-fold decrease in mean IL-2ra expression at the population level (Supplemental Fig. 1h-k & Supplemental Fig 2. i-l). This baseline deficiency was rescued following exposure to anti-CD3/CD28 antibodies; Tregs from CaRE3-EDEL animals upregulated IL-2ra to equivalent values compared to stimulated Tregs from WT or CaRE4-EDEL animals (Fig. 1b). These *in vitro* data collectively suggest that the CaRE4 enhancer selectively fine-tunes the expression kinetics of IL-2ra in activated Tconvs while the CaRE3 enhancer selectively fine-tunes the steady-state expression levels of IL-2ra in Tregs. These data further suggest that an excess of extracellular IL-2 can mitigate imbalances in the *IL2ra* expression parameters of activated Tconvs or Tregs, in line with our previous study focused on CaRE4 enhancer mutations(*29*). Although the enhancer deletions are germline, given the cell-type-restricted activity of each enhancer, we herein refer to the CaRE4 deletion as Tconv-EDEL and the CaRE3 deletion as Treg-EDEL for clarity.

The preceding data demonstrated how perturbing non-coding enhancers can modestly alter gene expression parameters in specific immune cell types. This behaviour stands in marked contrast to rare protein-coding perturbations, which operate ubiquitously and can exert pronounced functional effects on their gene targets (i.e., significant loss or gain of function). However, most genetic variants associated with common autoimmune disease risk or protection map to non-coding enhancers, not protein-coding regions(*13, 15, 16*). For instance, the non-coding SNP rs61839660 resides within the CaRE4 enhancer and appears to reduce the risk of developing Type 1 diabetes (T1D) (Odds Ratio = 0.62, p value = 2.8 x 10^-39^) yet increase the risk of developing Crohn’s Disease in humans(*34, 35*). This SNP mimics the effect of the Tconv-EDEL in activated Tconvs, albeit with a more modest effect on IL-2ra expression compared to the enhancer deletion(*29*). Similarly, the CaRE3 enhancer has been reported to harbor a candidate risk variant for human autoimmune thyroiditis(*13*). These observations raise a fundamental question: are small changes in cell-type selective IL-2RA expression parameters sufficient to alter disease risk at the organismal level(*29*)?

To investigate this question, we first studied the progression of autoimmune diabetes in WT NOD and Tconv-EDEL NOD mice. The NOD genetic background significantly increases the risk of developing spontaneous autoimmune diabetes with age and serves as an established model of human T1D(*36, 37*). To assess the incidence of disease over time, we monitored blood glucose levels weekly. These measurements revealed a striking phenotype: while the majority of WT NOD animals developed diabetes over the course of 30 weeks, Tconv-EDEL^+/+^ NOD animals were completely protected across all measured time points (Fig. 2a, d). Accordingly, Tconv-EDEL^+/+^ NOD mice displayed little to no immune cell infiltration in their pancreatic islets at 16-18 and 32-34 weeks of age compared to WT NOD controls (Supplemental Fig. 4a-d). Even Tconv-EDEL^+/-^ NOD animals displayed significant protection from developing diabetes over time and reduced immune cell infiltration in their pancreatic islets compared to controls (Fig. 2 b, d & Supplemental Fig. 4a-d). These findings are consistent with data suggesting that heterozygous variants can modulate the risk of developing human autoimmunity(*34, 38*). Thus, modest changes in IL-2ra expression kinetics due to non-coding variation can alter organismal disease risk.

**Figure 4.**
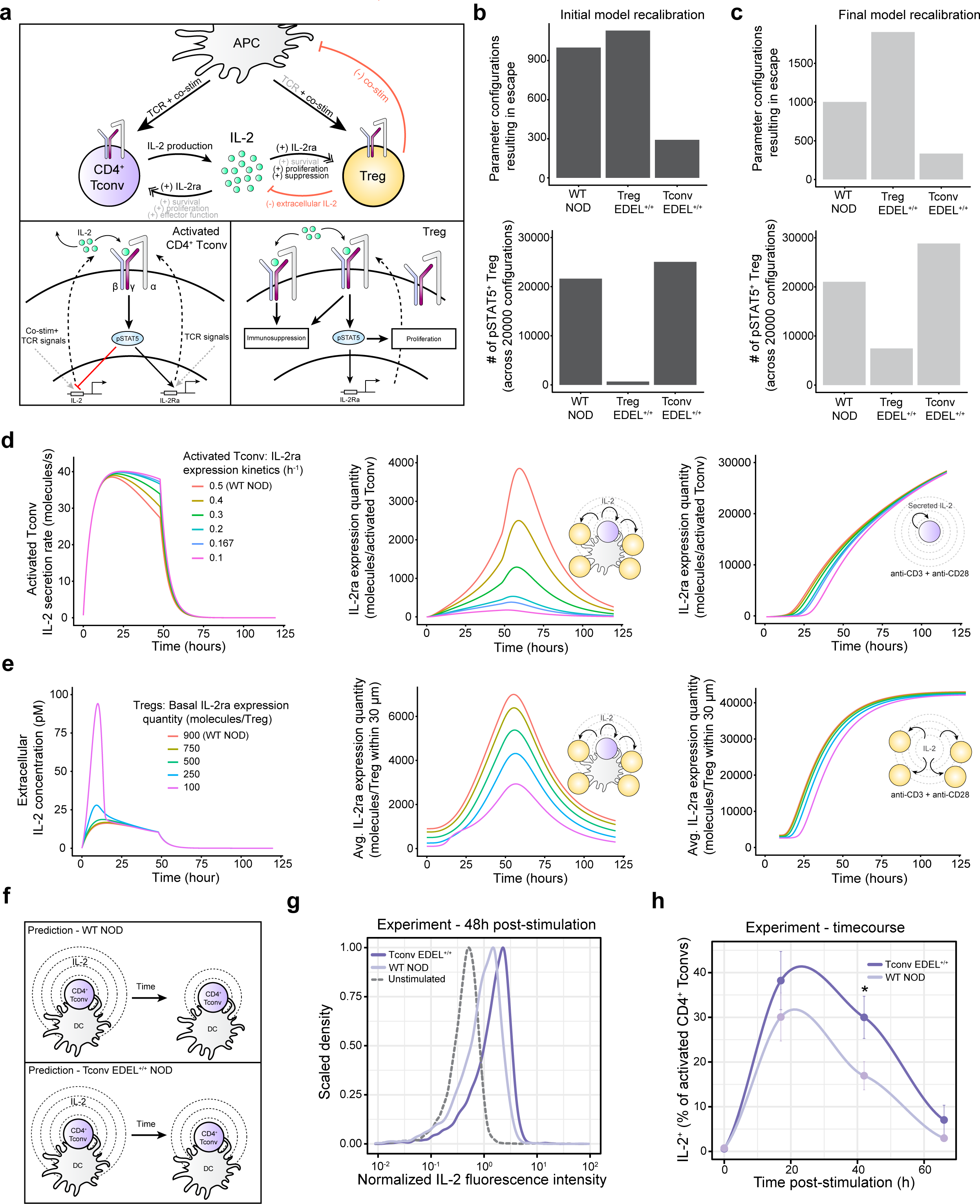
A multiscale computational model reveals how small changes in IL-2ra expression parameters alter intracellular signaling and intercellular communication dynamics within and between Tconvs and Tregs. a, Cartoon schematic illustrating three-cell multiscale model. Grey text = phenomenon not simulated by the model. Illustrated surface molecules represent the α, β, and γ chains of the IL-2 receptor **b and c,** summarized results from 20000 simulations using the initial and final recalibrated models for the NOD genotype, denoted as recalibration 1 and recalibration 3, respectively. Each simulation was initialized with a unique combination of biological plausible parameter values (i.e., parameter configurations). ‘Escape’ represents productive IL-2 sensing (i.e., a high pSTAT5 value) in the activated CD4^+^ Tconv during each simulation. **d,** dynamical trajectories of Tconv IL-2 secretion (left panel) and IL-2ra expression (middle and right panels) as the Tconv IL-2ra expression kinetics are decreased linearly, reflecting different perturbations to the CaRE4 enhancer. The WT NOD condition represents a parameter configuration that resulted in mild escape. Middle panel = Tconv IL-2ra expression dynamics following physiological stimulation *in vivo*. Right panel = Tconv IL-2ra expression dynamics following antibody-mediated stimulation and in the absence of APCs and Tregs. **e,** dynamical trajectories of extracellular IL-2 concentrations (left panel) and Treg IL-2ra expression (middle and right panels) as the baseline Treg IL-2ra is decreased linearly, reflecting a range of perturbations to the CaRE3 enhancer. The WT NOD condition represents the same parameter configuration as in d. Middle panel = Treg IL-2ra expression dynamics following physiological stimulation *in vivo*. Right panel = Treg IL-2ra expression dynamics following significant IL-2 levels *in vitro* in the absence of APCs. The amount of IL-2 equates that produced by a Tconv receiving strong antibody-mediated stimulation. **f,** Cartoon schematic illustrating an unexpected prediction from the computational model. **g,** Single-cell quantification of IL-2 production by purified CD4^+^ Tconvs with indicated genotypes. **h**, Pooled quantification of the percent of IL-2 producing CD4^+^ Tconvs at each indicated time point. Data are mean ± SEM per measured timepoint with n=3 animals from 2 independent experiments. Curves = local regression lines determined by the LOESS method. For g and h, CD4^+^ Tconvs were stimulated with low concentrations of anti-CD3/anti-CD28 (1μg/ml), with brefeldin A administrated 2.5h prior to each measured timepoint.

Several clinical immunotherapies target T cell functions to improve disease outcomes in human patients. This concept is best illustrated by checkpoint inhibitors, particularly anti-PD-1 blocking antibodies, which bolster the activation of Tconvs to eliminate tumours(*39, 40*). However, a significant fraction of treated patients experiences immune-related adverse events (irAEs) that resemble autoimmunity, particularly autoimmune diabetes(*41, 42*). Given the protective effect of the Tconv-EDEL on the development of spontaneous autoimmune diabetes, we hypothesized that it might also provide protection against checkpoint inhibitor-induced diabetes. To test this possibility, we administered anti-PD-1 blocking antibodies to WT NOD or Tconv-EDEL NOD animals and once again measured blood glucose levels over time. While anti-PD-1 blocking antibodies severely accelerated disease progression in WT NOD mice, consistent with previous findings, Tconv-EDEL^+/+^ NOD mice were completely protected from diabetes during the 30-day study with variable preservation of their pancreatic islets(*43*) (Fig. 2b, d & Supplemental Fig. 4e). Moreover, although Tconv-EDEL^+/-^ NOD animals still developed diabetes, they exhibited a significant delay in disease onset relative to WT NOD mice (Fig. 2b, d). These findings suggest that *IL2RA* enhancer variants with selective activity in stimulated Tconv cells may be predictive of irAEs in patients treated with checkpoint inhibitors.

Given the protective effects of the Tconv-EDEL on autoimmune diabetes, we hypothesized that changes in the IL-2ra expression parameters of Tregs would also have an impact on disease progression. Therefore, we assessed the incidence of spontaneous autoimmune diabetes in Treg-EDEL NOD animals compared to WT NOD controls. We found that 3-4 times as many Treg-EDEL^+/+^ NOD animals developed autoimmune diabetes at an accelerated pace compared to controls over the course of 30 weeks. More strikingly, perturbing the CaRE3 enhancer at a single *IL2ra* allele had an almost equivalent effect on disease progression (Fig. 2c,d & Supplemental Fig. 4f, g). These results suggest that a relatively small decrease in steady-state Treg IL-2ra expression can increase autoimmune disease risk markedly. These findings more generally highlight how the control of self-reactive Tconv responses by Tregs exhibits fragility with respect to IL-2ra expression variation, consistent with the fact that multiple autoimmune disease-associated variants map to non-coding sequences within the human *IL2RA* locus(*13*).

The marked and divergent disease outcomes observed in Tconv-EDEL vs. Treg-EDEL animals strongly suggested that the subtle, cell-type selective changes in *IL2ra* expression parameters we measured *in vitro* led to potentiated effects on immune regulation *in vivo*. We hypothesized that this potentiation may occur due to IL-2-dependent feedback circuits operating within and between activated Tconvs and Tregs, each with non-linear components that could skew IL-2 signalling towards one cell type or the other. For example, at the intracellular scale, IL-2ra expression can increase non-linearly due to positive feedback, which is only triggered once a Tconv or Treg senses a threshold amount of IL-2(*5, 44*). Furthermore, IL-2 signalling within activated Tconvs initiates negative feedback that can blunt the production of IL-2 itself(*45*). Lastly, at the intercellular scale, an IL-2-dependent spatial feedback circuit enables Tregs to constrain IL-2 production and sensing by weakly activated Tconvs in lymph nodes (LNs)(*3*). The multiscale operations of these circuits can likely amplify or dampen the destructive capacity of activated Tconvs in unexpected ways.

To examine how cell-type selective changes in IL-2ra regulation propagate *in vivo*, we employed quantitative multiplexed imaging to compare STAT5 phosphorylation (pSTAT5)—a readout of IL-2 signalling—in activated Tconvs and Tregs within the pancreatic LNs of WT and EDEL NOD mice (Fig. 3a & Supplemental Fig. 5a). We focused on CD4^+^ Tconvs because many enhancer variants associated with autoimmunity exhibit selective activity in this cell type compared to CD8^+^ Tconvs(*13, 20*). Within Treg-EDEL^+/+^ NOD animals, we observed a 3-fold increase in the frequency of pSTAT5^+^ CD4^+^ Tconvs and a 2-fold decrease in the frequency of pSTAT5^+^ Tregs compared to WT NOD controls. By contrast, within Tconv-EDEL^+/+^ animals, we observed inverse phenotypes, with a ∼3-fold decrease in pSTAT5^+^ CD4^+^ Tconvs and a ∼2.5-fold increase in pSTAT5^+^ Tregs (Fig. 3a & Supplemental Fig. 5a). In all animals, pSTAT5^+^ CD4^+^ Tconvs exhibited elevated expression of the inhibitory receptor PD1—a specific readout of productive TCR-mediated signalling—suggesting that these cells were activated by endogenous self-antigens displayed on antigen-presenting cells (APCs) within the pancreatic LNs(*3*) (Fig. 3b).

**Figure 5.**
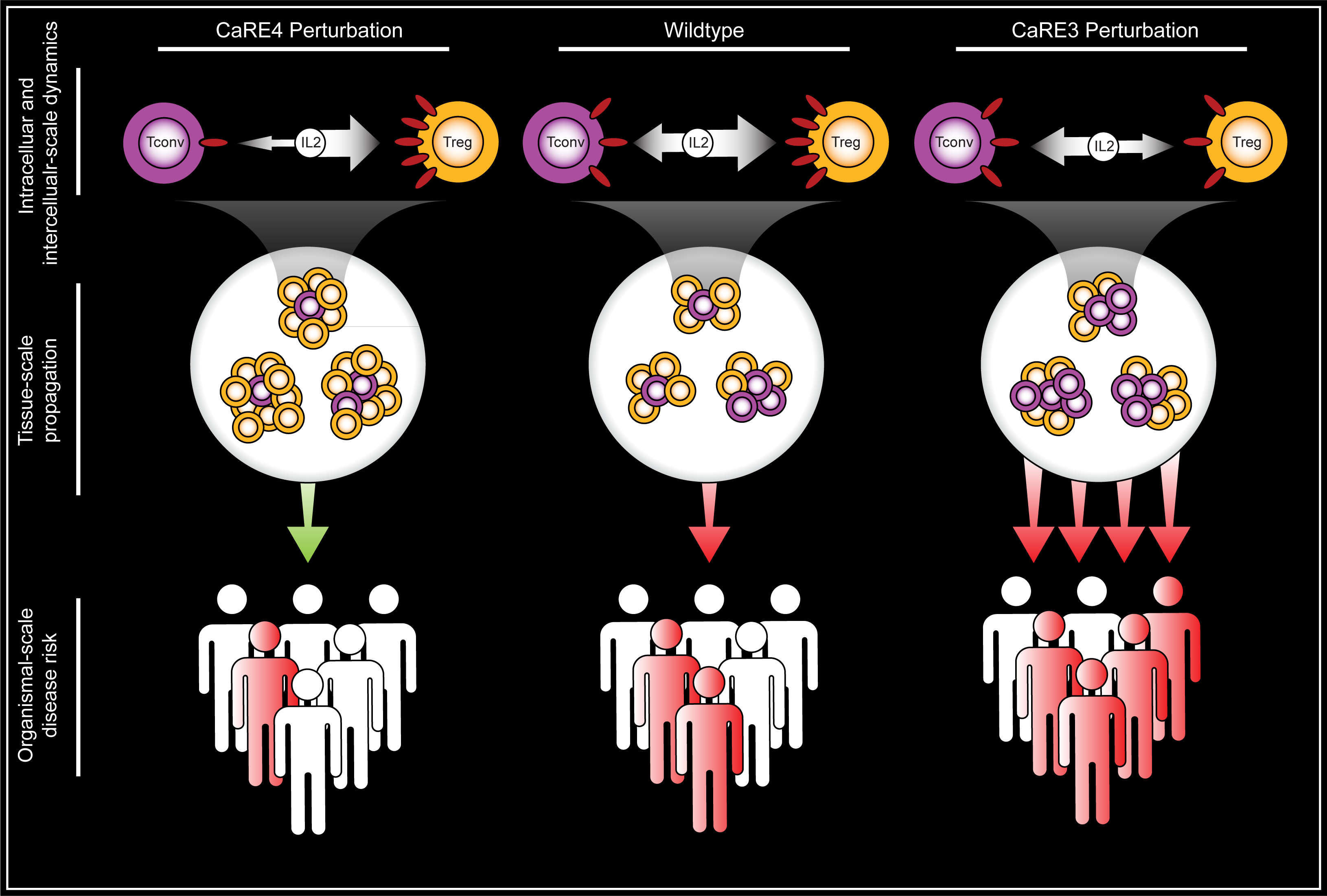
Conceptual model for how small changes in Tconv or Treg *IL2RA/Il2ra* expression parameters can propagate across biological scales to shape organismal disease risk.

The effects of each EDEL on the control of autoimmune Tconv responses propagated beyond the marked skewing of IL-2 signaling, causing dramatic spatial re-organization at the tissue scale. We observed changes in the local densities of Tregs or activated Tconvs throughout the pancreatic LN paracortex, resulting in opposing spatial domains that favoured the activity of one cell type or the other (Fig. 3c-e). This phenotype is consistent with the dichotomous capacity of IL-2 to modulate the immune response. In Tconv-EDEL^+/+^ NOD animals, the local density of Tregs increased significantly surrounding individually activated Tconvs. This increase coincided with a low IL-2Ra expression ratio between each activated Tconv and their most proximal Tregs (Fig. 3d, e & Supplemental Fig. 5 b-d). Thus, in this *in vivo* setting, activated Tconvs lacking the CaRE4 enhancer failed to develop high surface expression of IL-2ra. These results differed from our *in vitro* experiments in which activated Tconvs from Tconv-EDEL animals eventually expressed high levels of IL-2ra. This discrepancy is most likely explained by high IL-2 concentrations in the culture due to 1) prolonged Tconv stimulation by anti-CD3/CD28 antibodies, and 2) the inability of Tregs to constrain Tconv co-stimulation via CTLA-4—a critical immunosuppressive mechanism that is bypassed by anti-CD28 antibodies(*46–48*).

In contrast to the Tconv-EDEL^+/+^ NOD animals, we observed dense spatial domains of activated Tconvs in Treg-EDEL^+/+^ NOD mice, consistent with local bursts of proliferation and enhanced survival (Fig. 3c, d). These results suggested that in Treg-EDEL^+/+^ NOD mice, local Treg control was insufficient to constrain most incipient autoimmune responses, even though Tregs occasionally exhibited elevated densities within some of the activated Tconv clusters. In Treg-EDEL^+/+^ NOD animals, many activated Tconvs displayed high expression levels of IL-2Ra, and thus, a significantly elevated IL-2ra expression ratio between each activated Tconv and their proximal Tregs compared to WT or Tconv-EDEL^+/+^ NOD animals (Fig. 3d, e & Supplemental Fig. 5 b-d). These results once again differed from our *in vitro* experiments in which Tregs from Treg-EDEL^+/+^ animals eventually showed high surface expression of IL-2ra following strong antibody-mediated stimulation, most likely due to the high availability of IL-2 in the culture. Overall, these data demonstrate that small, cell type-specific changes in *IL2ra* expression parameters are potentiated *in vivo*, non-linearly boosting the competing functions of activated Tconvs or Tregs.

To gain quantitative, mechanistic insights into how this potentiation occurs, we simulated control of an autoimmune Tconv response within WT or EDEL NOD animals by modifying our previously described multiscale computational model, which was largely based on parameter ranges obtained from experiments using C57BL/6 mice(*3, 49*) (Fig. 4a). This modeling approach can provide critical insights into how variation in Tconv, Treg, or APC function affect the progression of an activated Tconv response. We simulated the activation of a CD4^+^ Tconv by a self-antigen-bearing APC, leading to the secretion of IL-2. We further simulated a spatially-resolved cluster of Tregs, which upon sensing secreted IL-2, increase in local density as well as immunosuppressive functionality, dampening both IL-2 production and autocrine sensing by the activated CD4^+^ Tconv. For an autoimmune response to progress, the activated Tconv must escape from these local Treg constraints and sense a threshold amount of IL-2, which is read out by measuring the maximum pSTAT5 signal achieved over the course of 120 hours *in silico*. This escape phenotype depends heavily on the operations of multiple IL-2-dependent feedback circuits—both at the intracellular and intercellular scales—which have been described previously in the literature and our own work(*3, 5, 28, 50*). Our model is governed by fixed parameters with static values (e.g., IL-2 transcriptional rate) and variable parameters with defined ranges (e.g., TCR off-rates for self-antigens, the local density of Tregs etc.). For individual simulations, fixed parameters are held constant while combinations of values for each variable parameter, known as parameter configurations, are sampled at random from biologically plausible ranges.

We initially recalibrated parameter ranges in our model to account for the distinct genetic background of NOD mice compared to C57BL/6 mice, with the overarching goal of recapitulating our experimental observations and subsequently making novel predictions that we could experimentally validate. To this end, we extended the lower range of TCR off-rates for self-antigens among CD4^+^ Tconvs, reflecting the fact that WT NOD animals harbour a unique MHC haplotype that alters the thymic selection process, skewing the Tconv repertoire towards increased self-reactivity relative to C57BL/6 mice(*51–54*) (Supplemental Fig. 6a). This recalibration failed to recapitulate our observed experimental observations, presumably because NOD mice also contain a range of non-MHC genetic variants that alter Tconv and Treg functions compared to C57BL/6 mice (Fig. 4b). Indeed, only a small subset of parameter configurations displayed an increase in CD4^+^ Tconv escape in response to the Treg-EDEL^+/+^ genotype relative to WT NOD mice, and only a small subset of parameter configurations displayed an increase in pSTAT5^+^ Tregs in response to the Tconv-EDEL^+/+^ genotype (Fig. 3a, Supplemental Fig. 5a, Supplemental Fig. 6a). These discrepancies between the simulations and our experimental findings prompted us to recalibrate the model further. First, we accounted for the fact that Tregs are intrinsically less functional in NOD versus C57BL/6 mice, expressing lower quantities of IL-2ra and Foxp3 for instance(*55*) (Supplemental Fig. 6b, c). Second, we employed a computational framework called “MAchine learning of Parameter-Phenotype Analysis” (MAPPA) to selectively filter our initial simulations for the few parameter configurations that matched our experimental results in Tconv-EDEL^+/+^ NOD animals (i.e., increased pSTAT5^+^ Tregs)(*49*). MAPPA then constructed a random forest machine learning model to assess the underlying parameter ranges in this subset of configurations compared to others (Supplemental Fig. 6d-f). Unexpectedly, the random forest model revealed lower values of the pSTAT5_EC50^IL-2^ parameter in the configurations that matched our experimental results, reflecting increased negative feedback on IL-2 production by pSTAT5 in the activated CD4^+^ Tconvs (Supplemental Fig. 6d-f). Based on these collective findings, we rationally adjusted a small number of parameters, both fixed and variable, and ran 20000 new simulations for the WT, Treg-EDEL^+/+^, and Tconv-EDEL^+/+^ NOD genotypes (Supplemental Fig. 6f). These new simulations were far more concordant with our experimental data, demonstrating that the recalibrated model better accounted for the genetic background of NOD mice (Fig. 4c).

Using this refined model for NOD mice, we proceeded to investigate how the cell-type-selective effects of the Tconv-EDEL and Treg-EDEL were potentiated *in vivo* by analyzing the simulated dynamic trajectories of IL-2 production, IL-2ra expression, and pSTAT5 signalling. This analysis produced several non-intuitive findings. Specifically, in the context of the Tconv-EDEL^+/+^ genotype, the model predicted an unexpected increase in the duration of IL-2 production by activated CD4^+^ Tconvs (Fig. 4d). To better explore this result, we linearly decreased the IL-2ra expression kinetics in the activated CD4^+^ Tconv from 0.5 h^-1^ (WT NOD) down to 0.1 h^-1^, and in all cases, we observed a corresponding linear increase in IL-2 production duration (Fig. 4d). These results indicate that lower IL-2ra expression is associated with reduced intracellular negative feedback on IL-2 production by IL-2-induced pSTAT5 (Supplemental Fig. 6d, e). In other words, kinetic delays in IL-2ra expression cause an activated CD4^+^ Tconv to sense less IL-2, resulting in diminished STAT5 phosphorylation, and therefore, increased IL-2 production duration. The model further predicted that such increased IL-2 production would enhance IL-2 sensing in surrounding Tregs, thereby enhancing their pSTAT5 signal, local density, and immunosuppressive function (Supplemental Fig. 7a). As a result, the activated CD4^+^ Tconv is subjected to an increasing amount of spatially localized, intercellular negative feedback at longer timescales, severely restricting its ability to sense IL-2. This lack of IL-2 signalling also limits the activated CD4^+^ Tconv from initiating an intracellular positive feedback circuit that non-linearly boosts IL-2ra expression—especially when the Tconv IL-2Ra expression rate falls below 0.3 h^-1^—further skewing IL-2 signals in favor of Tregs (Fig. 4d). These simulations provided a mechanistic explanation for three experimental observations in the pancreatic LNs of Tconv-EDEL^+/+^ NOD mice: 1) negligible pSTAT5 signal in activated CD4^+^ Tconvs, 2) increased pSTAT5 signal in Tregs, and 3) and elevated local densities of Tregs surrounding each activated CD4^+^ Tconv (Fig. 3a-e). Our model also revealed that *in vitro* stimulation conditions (i.e., anti-CD3/anti-CD28 antibodies without Tregs) promoted strong autocrine IL-2 sensing in the activated CD4^+^ Tconv, which in turn, induced positive feedback upregulation of IL-2ra (Fig. 4d). These *in silico* results demonstrated that in the absence of functional Tregs (or excess IL-2 as in the culture experiments), what is normally slow IL-2ra expression by activated CD4^+^ Tconvs is quickly negated over time. In the presence of Tregs and limiting IL-2 amounts, however, such decreases are significantly amplified and propagated across scales due to the compounded effects of multiple IL-2-dependent feedback circuits, significantly blunting the progression of an ongoing autoimmune response (Fig 3a-e, Fig. 4d, Supplemental Fig. 7a).

In the context of the Treg-EDEL^+/+^ genotype, a similar analysis of dynamic trajectories revealed critical insights into why autoimmune CD4^+^ Tconv responses were potentiated *in vivo*. Most importantly, the model predicted that the Treg-EDEL^+/+^ genotype would limit IL-2ra expression in Tregs while enhancing that in activated CD4^+^ Tconvs far beyond the level observed in WT NOD mice (Fig. 4e, Supplemental Fig. 7b). These outcomes were consistent with the locally skewed IL-2ra expression ratios we observed *in situ* (Fig. 3d, e). To gain further insights into this prediction, we linearly decreased the baseline Treg IL-2ra expression from 900 molecules/cell (WT NOD) down to 100 molecules/cell *in silico*. This titration resulted in non-linear IL-2ra dynamics, both in Tregs and the activated CD4^+^ Tconv, once the baseline Treg IL-2ra expression quantity fell to ∼250 molecules/cell. At or below this value, the local concentration of extracellular IL-2 increases dramatically due to insufficient IL-2 absorption by Tregs, thus enabling productive IL-2 sensing and STAT5 phosphorylation by the activated CD4^+^ Tconv (Fig. 4e, Supplemental Fig. 7b). This effect boosts the latter’s IL-2ra expression to promote further IL-2 capture, which should enhance cell proliferation and survival (Supplemental Fig. 7b). Interestingly, although the resulting increase in pSTAT5 signal initiates negative feedback on IL-2 production, the extracellular cytokine concentration stays within a range that sustains the activated CD4^+^ Tconv response but fails to promote a local Treg response (Supplemental Fig. 7b). These simulations explained why the local densities of activated CD4^+^ Tconvs were elevated in the pancreatic LNs of Treg-EDEL^+/+^ NOD mice (Fig. 3 c, d). We further simulated Treg stimulation *in vitro* in the absence of a competing CD4^+^ Tconv. In this scenario, Tregs effectively engaged IL-2 and rapidly upregulated high surface expression of IL-2ra over time due to IL-2-dependent positive feedback (Fig. 4e). Thus, baseline reductions in Treg IL-2ra expression, such as those observed in the Treg-EDEL^+/+^ or Treg-EDEL^+/-^ NOD animals, are negated in the absence of activated CD4^+^ Tconvs that compete for the cytokine. The presence of IL-2ra competent Tconvs, however, significantly potentiates early imbalances in Treg IL-2ra expression due to the multiscale operations of IL-2 feedback circuits, thereby promoting an ongoing autoimmune response (Fig. 3a-e, Fig. 4e, Supplemental Fig. 7b).

These computational studies suggested specific mechanistic explanations for how relatively small, T cell-selective changes in IL-2ra expression parameters could significantly alter the progression of autoimmune responses *in vivo* and shape organismal disease risk. The most unexpected of these was the increase in IL-2 production duration by activated CD4^+^ Tconvs with the Tconv-EDEL^+/+^ genotype due to reduced IL-2 sensing (Fig. 4d, f). To experimentally examine this key prediction, we purified naive CD4^+^ Tconvs from WT or Tconv-EDEL^+/+^ NOD mice and stimulated them *in vitro* using low rather than high concentrations of anti-CD3/anti-CD28 antibodies, better mimicking the physiological level of input signal these cells likely encounter *in vivo*. We then measured IL-2 production dynamics by flow cytometry. Because IL-2 is secreted rapidly upon synthesis, precluding its detection, we exposed activated CD4^+^ Tconvs to Brefeldin A for 2.5 hours prior to each measured time point(*56, 57*). This approach enabled short-term accumulation of IL-2 within the ER, ensuring a measurable quantity of cytokine per cell while maintaining relatively accurate temporal resolution (Figure 4g). As predicted by our computational model, we observed prolonged IL-2 production duration by activated CD4^+^ Tconvs from Tconv-EDEL^+/+^mice compared to WT NOD mice (Figure 4g, h). These findings highlight how sustained, subtle changes in extracellular IL-2 concentrations can have profound effects on disease progression, consistent with the fact that low dose IL-2 immunotherapy prevents WT NOD animals from developing autoimmune diabetes(*58*). Furthermore, this combination of simulation and experiment provides important insights into how *IL2ra* variants can modulate IL-2 production in addition to IL-2 sensing.

Although genome-wide association studies (GWAS) and fine-mapping studies have identified numerous genetic variants associated with autoimmune diseases in non-coding control regions—especially those operating in T cell subsets—the mechanisms by which these variants meaningfully contribute to disease risk have remained unclear. Our combination of enhancer perturbations, multiplexed tissue imaging, and computational modelling shed light on this question by demonstrating how relatively subtle yet sustained changes in cell-type-selective gene regulation can propagate across biological scales, significantly shifting autoimmune disease susceptibility at the organismal level (Fig. 5). Collectively, these findings provide a framework for prioritizing and interpreting autoimmune disease-associated variants and, more generally, linking quantitative changes in gene expression to organismal phenotypes.

Our findings are consistent with prior work documenting how the local Treg density in LNs controls the integration of multiple signals required by activated Tconvs to sustain their proliferative burst, survive long-term, and adopt tissue-damaging effector functions. Because the local Treg density is adjusted dynamically through an intercellular IL-2-dependent circuit, which itself contains nested feedbacks operating at distinct spatiotemporal scales, relatively small changes in the production or sensing of extracellular IL-2 rapidly can skew an immune response in favour of activated Tconvs or Tregs(*3*). Such multiscale coupling explains why cell type-selective imbalances in IL-2RA expression parameters can alter the progression of autoimmunity. While the effects of the CaRE3 and CaRE4 EDELs on *IL2RA*/*IL2ra* expression parameters (within ∼1.5-3 fold) may be accentuated compared to those caused by naturally-occurring disease-associated human SNPs, our computational modelling approaches emphasize that even smaller changes are sufficient to cause non-linear breakdowns in the control of self-reactive Tconv responses. In support of this concept, our experiments demonstrated that heterozygous EDELs had significant effects on organismal disease risk, especially at the CaRE3 region. These data are consistent with the fact that non-coding SNPs in the *IL2RA* locus are linked to at least 8 human autoimmune disorders(*13*). Several non-coding variants associated with human autoimmunity also map to the *IL2* locus itself or loci associated with the IL-2 signaling machinery(*59–63*).

The conceptual findings in this study likely generalize to loci beyond *IL2RA*. Indeed, Tconvs and Tregs employ numerous control circuits independent of IL-2, operating from the earliest stages of TCR-mediated signal transduction to later stages of cytokine-mediated communication(*1, 2, 4, 64*). Such hierarchical control gives rise to sharp functional boundaries (i.e., checkpoints) that constrain or amplify ongoing immune responses at different points in time and space. This design principle ensures that individual cells or group of cells commit to qualitatively discrete states following defined input signals. Thus, quantitative shifts in the positioning or steepness of functional boundaries can markedly alter a system’s propensity to adopt specific states, and thus, scale Tconv responses(*3, 11*). We suggest that such shifts emerge physiologically from uncompensated variation in fragile circuit parameters, resulting in altered disease risk(*8, 11, 12*). This view is consistent with the fact that many non-coding variants associated with autoimmune diseases map to loci underlying known T cell feedback machinery(*20*). These findings have important implications for predicting the emergence of host autoimmunity, both spontaneous and iatrogenic forms.

More generally, our data demonstrate how the compounded intracellular, intercellular, and spatial behaviours of multiple cell types produce unique output response dynamics compared to single cells in isolation. These findings emphasize the need for integrative, multicellular frameworks to determine correctly how variation in gene expression parameters alters the progression of immune responses *in vivo*. We anticipate that such frameworks will be essential for interpreting modern disease-association studies and defining non-coding mechanisms of disease.

## ACKNOWLEDGEMENTS

We thank all members of the Marson, Germain, and Wong labs. We thank Jeffrey A. Bluestone for his mentorship and guidance, and we thank Stacie Dodgson for critical reading of the manuscript. D.R.S. was supported by the Jeffrey G. Klein Family Fellowship in Diabetes. J.T.C. was supported by the National Science Foundation Graduate Research Fellowship Program. This was work supported funds from NIH/NIDDK (DP3DK111914-01), the Chan Zuckerberg Biohub, the Northern California JDRF Center of Excellence, the Larry L. Hillblom Foundation (grant no. 2020-D-002-NET), the Intramural Research Program of the National Institute of Allergy and Infectious Diseases, NIH, and the Ragon Institute of MGH, MIT and Harvard. The Marson lab has received gifts from J. Aronov, G. Hoskin, K. Jordan, B. Bakar, the Caufield family and funds from the Innovative Genomics Institute (IGI), and the Parker Institute for Cancer Immunotherapy (PICI). A.M. holds a Career Award for Medical Scientists from the Burroughs Wellcome Fund, is an investigator at the Chan–Zuckerberg Biohub and is a recipient of a The Cancer Research Institute (CRI) Lloyd J. Old STAR grant. This work used the UCSF Flow Cytometry Core, supported by the Diabetes Research Center grants NIH P30 DK063720 and NIH S10 1S10OD021822-01.

## AUTHOR CONTRIBUTIONS

Conceptualization, D.R.S., M.S.A., J.A.B., R.N.G., H.S.W., A.M.; Methodology, D.R.S., K.P., H.S.W.; Investigation, D.R.S., J.T.C., Z.L., A.Y., K.P., J.U., A.C.I., J.M.W., H.S.W.; Resources, V.N., J.M.W.; Formal Analysis, D.R.S., K.P., H.S.W.; Supervision, M.S.A., R.N.G., H.S.W., A.M.; Funding Acquisition, R.N.G., H.S.W., A.M.; Writing – Original Draft, D.R.S., J.T.C., R.N.G., A.M., H.S.W.; Writing – Review & Editing, D.R.S., K.P., J.T.C., J.S.T., R.N.G., A.M., H.S.W.

## METHODS

### Mouse Generation

CaRE3 enhancer deletion mice were generated at UCSF by electroporating Cas9 RNPs into mouse blastocysts(*66*). Briefly, equal volumes of 160uM CaRE3 1-4 crRNA (IDT) were mixed. An equal volume of 160uM tracrRNA was then added and the gRNA components were allowed to hybridize by incubating at 37C for 10 minutes. We then added an equal volume of 40uM Cas9 protein (QB3 Macrolab) and incubated the solution at 37C for 10 minutes to allow the Cas9 RNP to complex. Mouse lines were established by backcrossing founders to wild type animals at least one generation before performing experiments. Genotyping was done by PCR to look for the expected enhancer deletion. In total four founder lines were established, one on the NOD/ShiLtJ. background and three on the C57BL/6J background. All founders were immunophenotyped and phenotypes were found to be consistent. CaRE4 enhancer deletion mice were generated by the Jackson Laboratory and were reported previously(*29*)

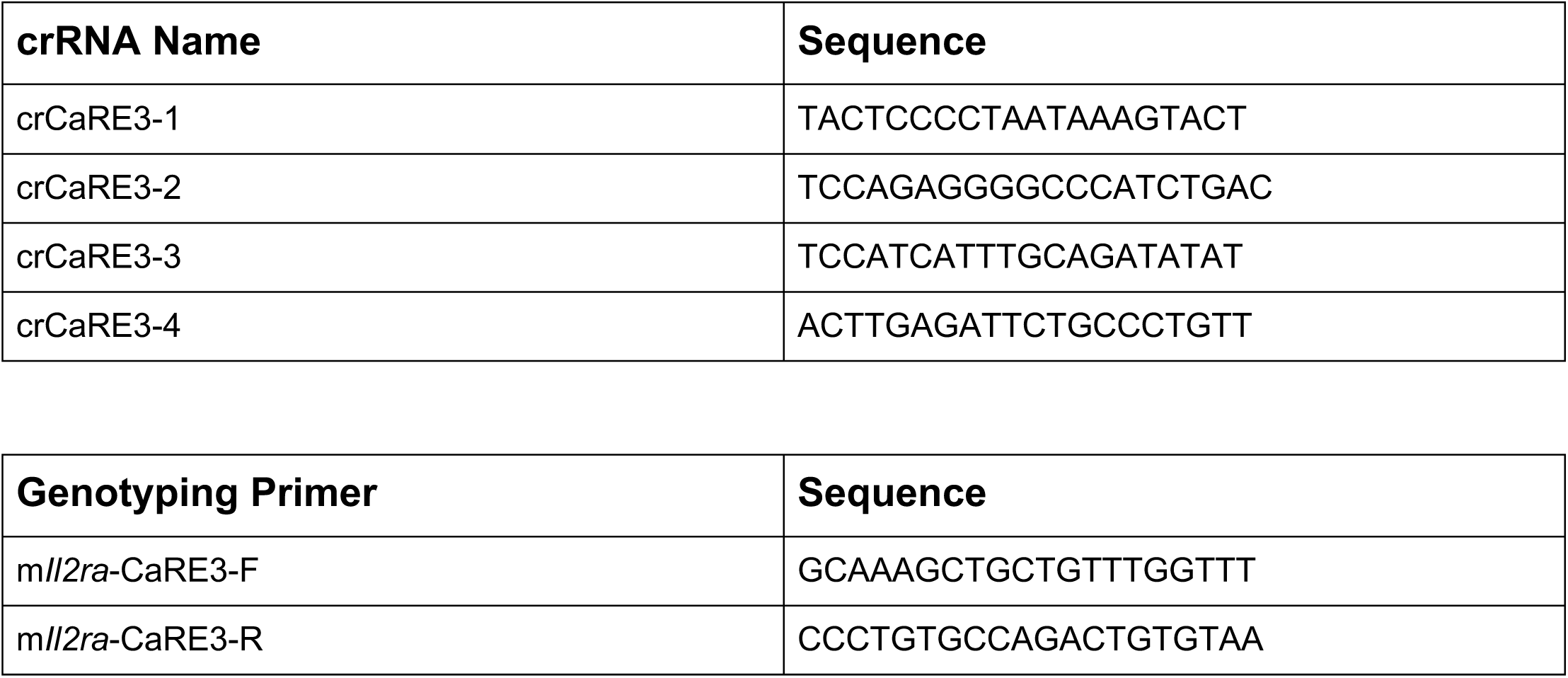

### Mouse Experiments and Data Analysis

All mice were maintained in the UCSF-specific pathogen-free animal facility in accordance with guidelines established by the Institutional Animal Care and Use Committee and Laboratory Animal Resource Center. Experiments were done with animals aged between 1 and 4 months. Wildtype littermate mice were used as controls for all immunophenotyping experiments. All mice used for experiments were normoglycemic unless otherwise noted. Data were excluded only for experimental reasons, either due to failed controls or insufficient numbers for meaningful quantification. Power calculations were not performed and data were assumed to be normally distributed. Experiments were done without blinding or randomization. All data are derived from at least two independent experiments unless otherwise stated. Both male and female NOD mice were used for immunophenotyping experiments.

During the course of evaluating the CaRE3 EDEL littermate control cohort for diabetes, tropical rat mites infected animals in our facility, requiring us to treat and quarantine our colony. Mice were treated with Permethrin nestlets for 6 weeks. The nestlet contains 7.4% of the active ingredient Permethrin which delivers 1800 mg of Permethrin per kg bodyweight, an amount that has been shown to be non-toxic to rodents. Nestlets (Ancare) were autoclaved before treatment was applied.

### Mouse Immunophenotyping Staining

Staining protocol of mouse cells was previously described(*29*). In short, staining was done in V-bottom 96 well plates. Surface staining was performed for 10-30 minutes on ice. We followed manufacturer’s protocols for fixation, typically 30 minutes at room temperature. Intracellular staining was performed for 30-60 minutes at room temperature. Antibodies were used at 1:100 dilution. The following staining panels were used in this study:

Thymus

*Surface and Viability*

Live/Dead-UVB (Invitrogen)

CD45-BUV395 (30-F11)

CD8-PerCP-Cy5.5 (53-6.7)

CD4-BV605 (RM4-5)

Il-2ra-PE-Cy7 (PC61.5)

CD69-APC (H1.2F3)

TCR beta-PB (H57-597)*

CD73-PE*

*Intracellular:*

Foxp3-FITC (FJK-16s)

\**not always included*

Spleen and inguinal LN:

*Surface and Viability:*

Live/Dead-UVB (Invitrogen)

CD45-BUV395 (30-F11)

CD8-PerCP-Cy5.5 (53-6.7)

CD4-BV605 (RM4-5)

Il-2ra-PE-Cy7 (PC61.5)

CD69-APC (H1.2F3)

*Intracellular:*

Foxp3-FITC (FJK-16s)

For experiments that required culture of cells complete RPMI with 10% FBS, HEPES, sodium pyruvate, non-essential amino acids, penicillin streptomycin, and beta-mercaptoethanol was used. For Il-2ra time course naïve CD4^+^ T cells were enriched from splenocytes using EasySep kit (StemCell Technologies). Approximately 100,000 cells were activated using 2ug/ml plate bound anti-CD3/CD28. Cells were stimulated for three days and surface Il-2ra expression was measured by flow cytometry every day.

### Splenocyte Stimulations

Splenocytes were plated at approximately 4 million cells per 500ul of complete RPMI in one well of a 48 well plate. Cells were activated using 1ug/ml anti-CD3 (Clone: 145-2C11, Biolegend) and anti-CD28 (Clone: 37.51, Biolegend). Cells were stained and analyzed by flow cytometry at the end of the time course using the following antibody panels:

#### Antibodies

Live/Dead-UVB (Invitrogen, Cat#: L23105)

CD45-BUV395 (30-F11)

TCR beta-BV421 (H57-597)

Il-2ra-PE-Cy7 (PC61.5)

CD4-BV605 (RM4-5)

CD8-PerCP-Cy5.5 (53-6.7)

Foxp3-FITC (FJK-16s)

### Spontaneous Diabetes

Litters of female NOD mice from CaRE3 and CaRE4 mice were set aside for spontaneous diabetes studies. Blood glucose was measured once a week from a drop of blood obtained from the end of the tail. We measured blood glucose in animals starting at 8-11 weeks of age until 30 weeks of age. Once animals entered the study they were not removed for any reason. The diabetes endpoint was defined as two consecutive measurements of blood glucose over 200mmol/L. Diabetic animals were euthanized according to institutional standards.

### Anti-PD1 Induced Diabetes

For this study we followed a previously published dosing regimen(*43*). Briefly, anti-PD1 (Clone: RMP1-14, BioXCell, Cat: BE0146) was administered intraperitoneally to 8-10 week old female NOD mice. Animals were initially dosed with 500ug of antibody (day 0), and then an additional five injections of 250ug each were given every other day (days 2, 4, 6, 8, 10). Mice were followed for a total of 30 days. The diabetes endpoint was defined as two consecutive measurements of blood glucose over 200mmol/L. Diabetic animals were euthanized according to institutional standards.

### Pancreas Histology for Immune Infiltration

Briefly, the pancreas was isolated and placed into zinc formaldehyde fixative overnight at 4C. The next day the tissue was washed with water, 5 minutes per wash with gentle rocking. Three more washes were done with 1x PBS in a similar manner. Tissue was transferred to 70% EtOH and stored at 4C before further processing. Pancreas processing from 32-week-old animals was done by the UCSF Histology Core (Diabetes Center, UCSF). H&E stained sections were mounted on slides and scored under a microscope. Pancreas processing for 16-week-old animals was done by HistoWiz which completed embedding, sectioning, H&E staining, and imaging. Pancreatic islets were categorized as being normal, or as peri-insulitis or insulitis depending on severity and overall appearance of immune infiltration. Examples are shown in Extended Data Fig. 4.

### Tissue section preparation, processing and immunostaining

pLNs were isolated from 6-week old female NOD mice and fixed for 14h at 4°C in BD Cytoperm/Cytofix (BD Bioscience, Cat#: 554722) diluted 1/4 in PBS. pLNs were washed 3x in PBS (10 min per wash), carefully trimmed of fat using a stereo dissection microscope and fine forceps, and dehydrated for 24 h in a 30% sucrose solution made in 0.1 M phosphate buffer. LNs were then embedded in optimal cutting temperature (O.C.T.) compound (Sakura Finetek, Cat#: 50-363-579), frozen on dry ice, and stored at -80°C. 18-50 μm sagittal LN sections were prepared using a cryostat (Leica) equipped with a 40 Surgipath DB80LX blade (Leica, Cat#: 14035843497). Cryochamber and Specimen cooling was set to -17°C.

Tissue sections were adhered to Superfrost Plus microscopy slides (VWR, Cat#: 48311-703), permeabilized using 0.1% Triton X-100 for 10 min at 22°C, blocked in 5% mouse serum for 1 h at 22°C, and washed in PBS. Tissue sections were next incubated with directly conjugated primary antibodies diluted in PBS for 15 h at 4°C. After washing 3x in PBS (10 min per wash) at 22°C, samples were mounted in Fluoromount-G (SouthernBiotech, Cat#: 0100-01), which was allowed to cure for a minimum of 14h at 22°C. All imaging was performed using No. 1.5 coverglass (VWR, Cat#: 48393-241). The following directly-conjugated antibodies were used for immunostaining: anti-CD3 (clone 17A2, 1 μg/ml), anti-CD4 (clone RM4-5, 1 μg/ml), anti-Foxp3 (clone FJK-16s, 1 μg/ml), anti-Il-2ra (clone PC61, 1 μg/ml), anti-PD1 (clone 29F.1A12, 1 μg/ml). Combinations of the following organic fluorophores were used: Brilliant Violet 421, Brilliant Violet 480, Alexa Fluor 555, eFluor 570, Alexa Fluor 594, eFluor660, Alexa Fluor 647, Alexa Fluor 488, and Alexa Fluor 700.

For pSTAT5 immunostaining, fixed tissue sections were permeabilized in pre-chilled 100% methanol for 18 min at -20°C, washed extensively in PBS, blocked in 5% donkey serum for 1 h at 22°C, and washed further in PBS. Tissue sections were next incubated with unconjugated anti-pStat5 (clone C11C5, 1.715 μg/ml) diluted in PBS for 15 h at 4°C. Following washing in PBS at 22°C, tissue sections were incubated with either Alexa 488, Alexa 594, or Alexa 647-conjugated donkey anti-rabbit F(ab’)2 fragments (0.483 μg/ml, Jackson ImmunoResearch Laboratories, Inc, Cat#: 711-546-152 or 711-6060152) for 2h at 22°C. Sections were then washed 4x in PBS (10 min per wash) at 22°C, stained with directly-conjugated antibodies as described above, and mounted in Fluoromount-G.

### Laser scanning confocal microscopy

Digital images were acquired using an upright Leica TCS SP8 X spectral detection system (Leica) equipped with a pulsed white light laser, 4 Gallium-Arsenide Phosphide (GaAsP) Hybrid Detectors (HyDs), 1 photomultiplier tube (PMT), 40x (NA = 1.3) and a motorized stage. For tissue sections (18-20 μm), images were acquired with a pixel size of 0.271-0.286 μm, z step size of 0.3-1 μm, and detector bit-depth of 12.

### Image processing and segmentation

Image files generated in LAS X software were converted into “.ims” files in Imaris software (Bitplane) and subjected to a 1 pixel Gaussian filter to reduce noise.

Segmentation of densely packed T cells was performed using a protocol modified from Li et al. and described in Wong et al.(*3, 67*). In brief, this process involved creating artificial T cell nuclei, which in turn were subjected to segmentation algorithms. Initially, “.ims” files were imported into Fiji(*68*). The brightness and contrast of the CD4 channel was adjusted linearly to thinly demarcate the plasma membranes of T cells. The adjusted CD4 channel was then converted into an 8-bit format, duplicated and inverted to create an *inverted sum* channel. Next, the original channel was subtracted from the *inverted sum* channel, producing a *high-contrast inverted* channel, which was subsequently binarized using the “Auto Local Thresholding” tool. Binarized images were then despeckled to remove noise and subtracted from the *inverted sum* channel to improve separation between artificial T cell nuclei. The final product was exported as a “.TIFF” image sequence, imported into the original “.ims” file in Imaris, and subjected to the “Surface Object Creation” module. Segmentation artefacts were excluded using a combination of sphericity and volume thresholds, as well as manual correction.

### Recalibration of Multiscale T cell Activation Model, MAPPA simulation, and quantification of phenotypes

Dynamical modeling of CD4^+^ Tconv activation by an APC along with surrounding Tregs constraints was performed as previously described(*3*). Several fixed and variable parameters were adjusted to account for the unique genetic background of NOD mice based on experimental observations in this study or in the literature (see Supplemental Table 1)(*69–71*). These adjustments included extending the lower limit of TCR off-rates (recalibration 1), decreasing the basal expression level of Treg IL-2ra to (recalibration 2), and decreasing the range of Treg contact efficiency as well as the half maximum level of pSTAT5 for suppressing IL-2 expression (recalibration 3).

The MAPPA framework was implemented as previously described(*3, 49*). In brief, MAPPA employs Random Forests to construct quantitative maps between parameter space and phenotypic space for complicated dynamical models that lack analytical solutions. This framework ultimately quantifies how individual parameters sampled throughout the plausible parameter space affect specific phenotypes.

We readjusted ranges for variable parameters and 20000 randomly sampled parameter configurations to account for each recalibration scheme, as outlined above. Using these sampled parameter configurations, we simulated the model for WT NOD, CaRE3 EDEL NOD, and CaRE4 EDEL NOD conditions over the course of 0 to 120 hours using MATLAB software(*3*). We adjusted additional parameters to study the effects of EDELs on NOD animals. For the CaRE3 EDEL^+/+^, we decreased the basal transcriptional rate and initial condition of Treg IL-2ra by 10-fold whereas for the CaRE4 EDEL, we reduced the Tconv TCR and CD28 regulated transcriptional rate of IL-2ra by 3-fold.

For model recalibrations 1 and 2, only a small fraction of the parameter configurations produced simulations that matched our experimental observations, specifically increased pSTAT5^+^ Treg frequencies in CaRE4 EDEL^+/+^ animals compared to WT NOD controls. We recalibrated the model further by using this limited set of configurations to train a new random forest model that predicted the observed fractional changes between WT NOD and CaRE4 EDEL^+/+^ conditions in terms of average Treg pSTAT5 signal within 20 !" of an activated CD4^+^ Tconv. We then computed “variable importance” metrics for each parameter to determine its contribution to this phenotype (Supplementary Fig. 6. d & e).

To better examine the dynamical trajectories of the model in detail, we selected a representative parameter configuration that exhibited substantial changes in the maximum pSTAT5 levels between WT NOD and EDEL NOD conditions. We then simulated gradual changes in parameter values, reflecting varying degrees of perturbations to the CaRE4 or CaRE3 enhancers. For the CaRE4 enhancer, the TCR and CD28 regulated transcription rate of Tconv IL-2ra was linearly titrated from 0.5 h^-1^ to 0.1 h^-1^. For the CaRE3 enhancer, the basal level of Treg IL-2ra was linearly titrated from 900 molecules/cell to 100 molecules/cell.

For each simulated time trajectory, we used the maximum pSTAT5 level as a phenotype of interest. Tconvs were considered pSTAT5^+^ if they fell within the top 5 percentile of maximum pSTAT5 across all 20000 parameter configurations. For Tregs, we implemented a different approach because these cells were simulated using partial differential equations describing local Treg densities and intracellular states. As a result, we quantified number of pSTAT5^+^ Tregs using a 3 step process involving 1) obtaining the average maximum pSTAT5 level of Tregs within varying distances from the CD4^+^ Tconv and APC (from 10μm to 100μm), 2) determining the distance with the maximum pSTAT5 above the threshold value, and 3) integrating the Treg density function within that distance to determine the number of pSTAT5^+^ Tregs.

### Statistics

Statistical calculations were performed using GraphPad prism versions 7.0 and 8.0 or in the R statistical environment. Tests between two groups used a two-tailed Student’s t-test. Tests between multiple groups comparing EDEL to WT littermate controls used two-way ANOVA with Sidak’s test for multiple comparisons.

### Data availability

All data generated or analyzed are included in the published Article and the Supplementary Information. GWAS SNP tracks in Figure 1a derived from papers referenced in figure legend. H3K27Ac ChIP-seq data in Figure 1b for Tnaive (E039 Primary T helper naïve cells from peripheral blood), Tstim (E041 Primary T helper cells PMA-I stimulated), and Treg (E044 Primary T regulatory cells from peripheral blood) was generated as part of the Epigenome Roadmap Project (https://egg2.wustl.edu/roadmap/web_portal/). The FOXP3 and STAT5 ChIP-seq data in Figure 1b is publicly available as part of the FANTOM5 project. Data was uploaded to UCSC Browser for visualization and export (http://www.ag-rehli.de/TrackHubs/hub_Tsub.txt). There are no restrictions on data availability.

---

**Supplementary Table 1.** Ranges of parameter values used for the MAPPA framework

| **Name** | **Name in the main text** | **min** | **max** |
| --- | --- | --- | --- |
| $\nu$ | TCR<sup>off-rate</sup> | 0.2 (0.15 for R1) | 0.4 |
| $K_{TCR \rightarrow IL2R\alpha}$ | TCR_EC50<sup>IL-2Rα</sup> | 0.1 | 1 |
| $K_{costim \rightarrow IL2R\alpha}$ | CD28_EC50<sup>IL-2Rα</sup> | 10 | 100 |
| $K_{JAK \rightarrow pSTAT5}$ | JAK_EC50<sup>pSTAT5</sup> | 0.01 | 0.1 |
| $K_{TCR \rightarrow IL2}$ | TCR_EC50<sup>IL-2</sup> | 0.1 | 1 |
| $K_{costim \rightarrow IL2}$ | CD28_EC50<sup>IL-2</sup> | 10 | 100 |
| $K_{pSTAT5 \rightarrow IL2}$ | pSTAT5_EC50<sup>IL-2</sup> | 0.1 (0.05 for R3) | 1 (0.5 for R3) |
| $D_{IL2}$ | IL-2<sup>D</sup> | 10 | 100 |
| $n_{tr}$ | Treg<sup>λ</sup> | 0.0001 | 0.001 |
| $f_{cont\_low}$ | Treg<sup>Contact_baseline</sup> | 0.1 (0.075 for R3) | 0.4 (0.3 for R3) |
| $f_{cont\_high}$ | Treg<sup>Contact_stimulated</sup> | 1 | 2.5 |
| $L_{antigen}$ | [Antigen] | 100 | 1,000 |
| $L_{CD80\vert CD86}$ | [CD80/86] | 100,000 | 1000,000 |

R1, recalibration 1; R3, recalibration 3

**Supplementary Fig. 1.**
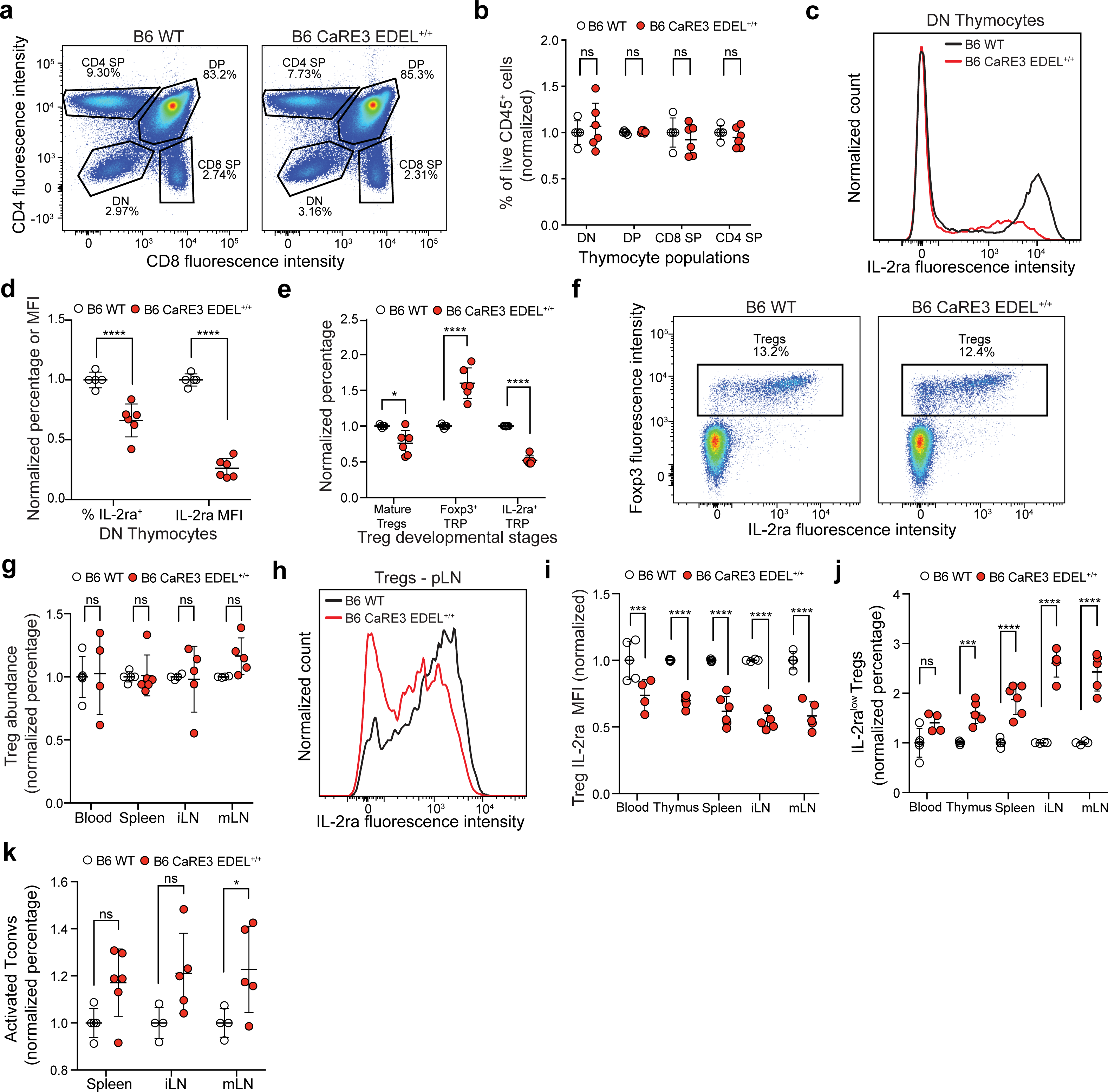

**Supplementary Fig. 2.**
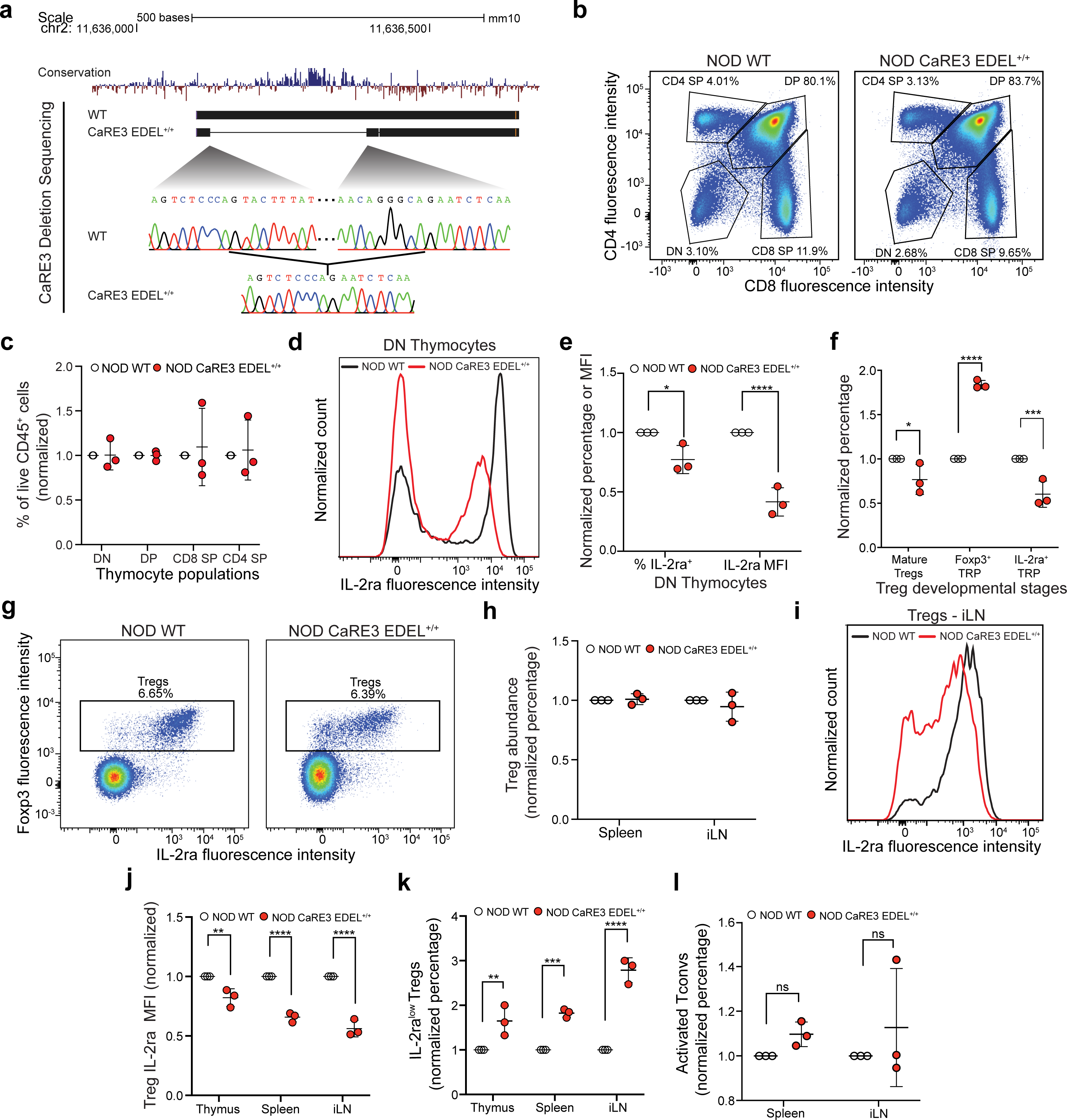

**Supplementary Fig. 3.**
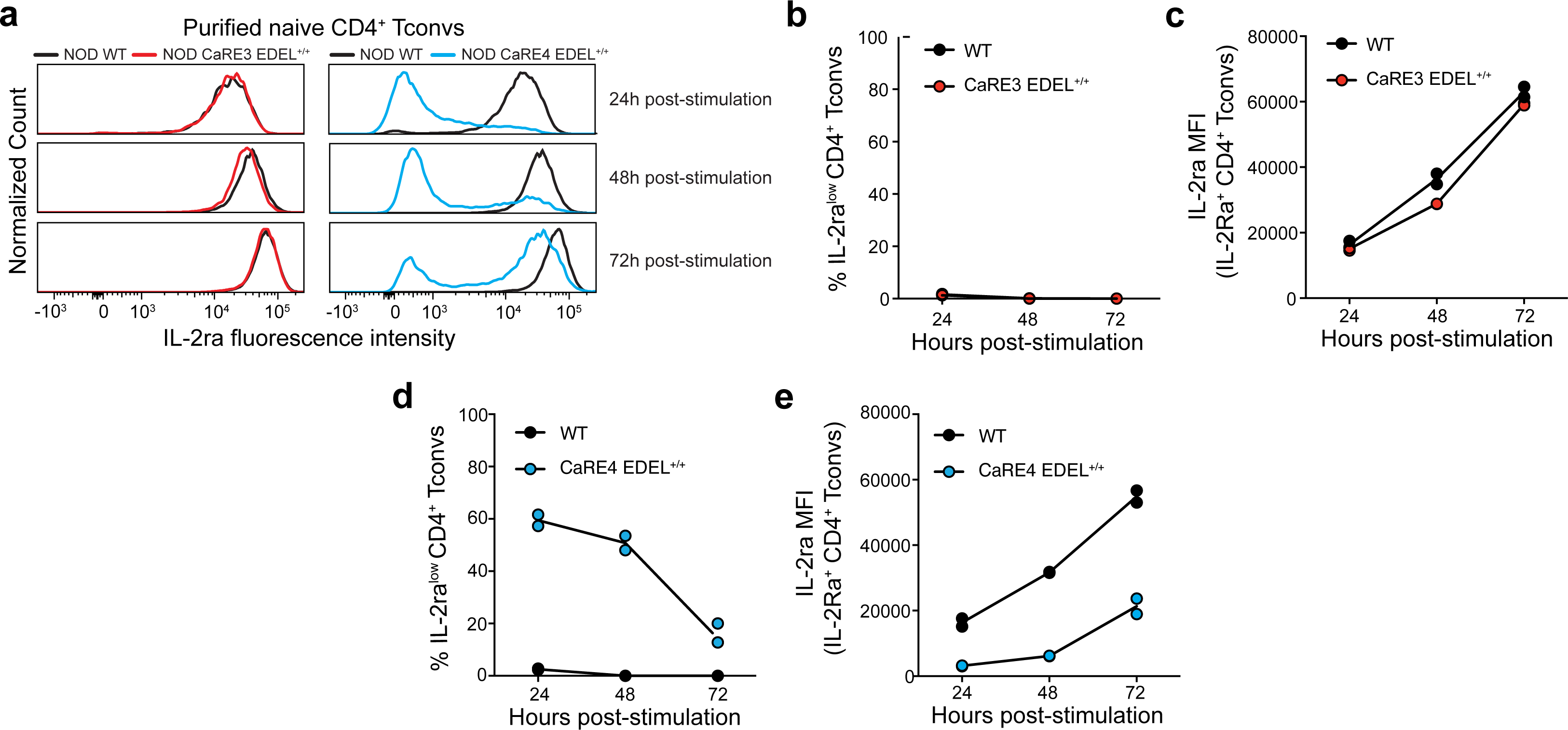

**Supplementary Fig. 4.**
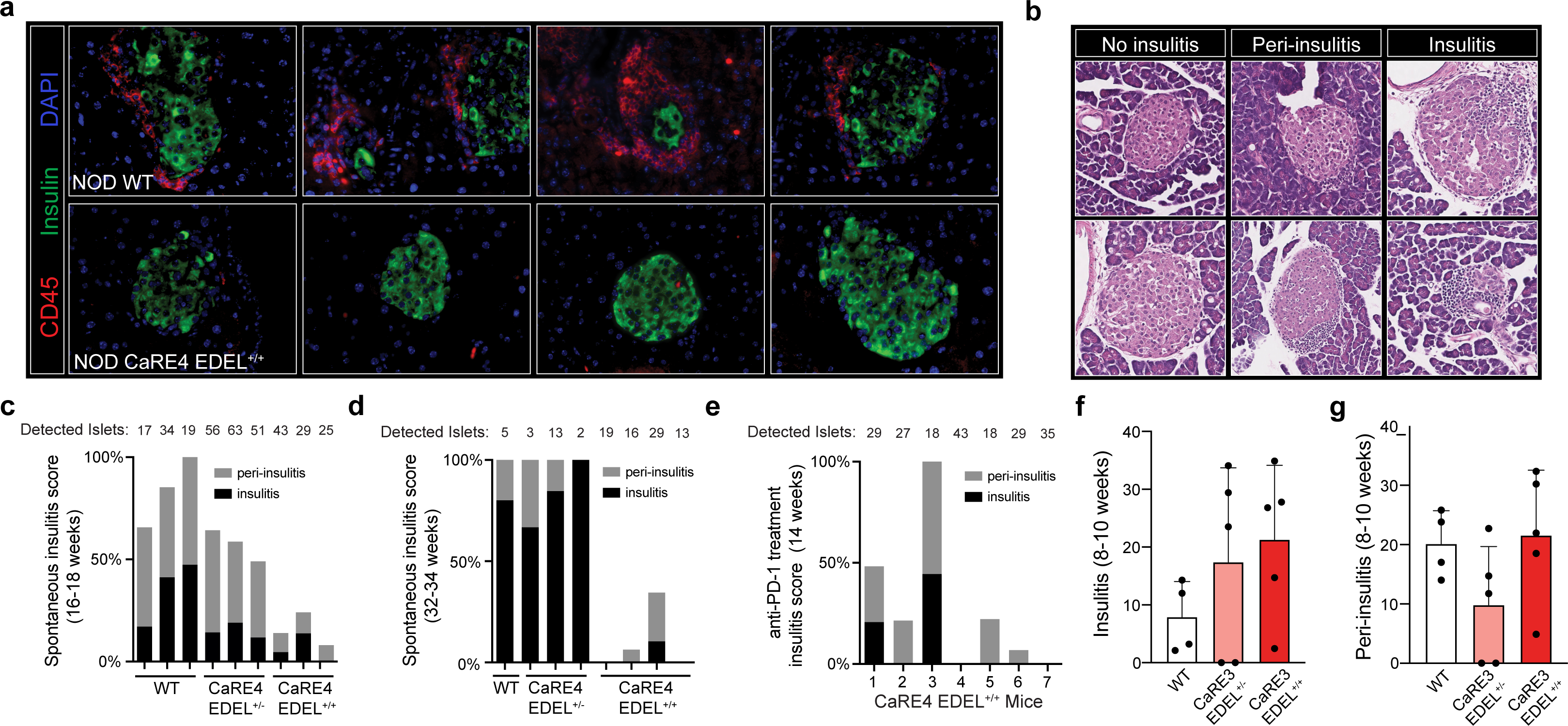

**Supplementary Fig. 5.**
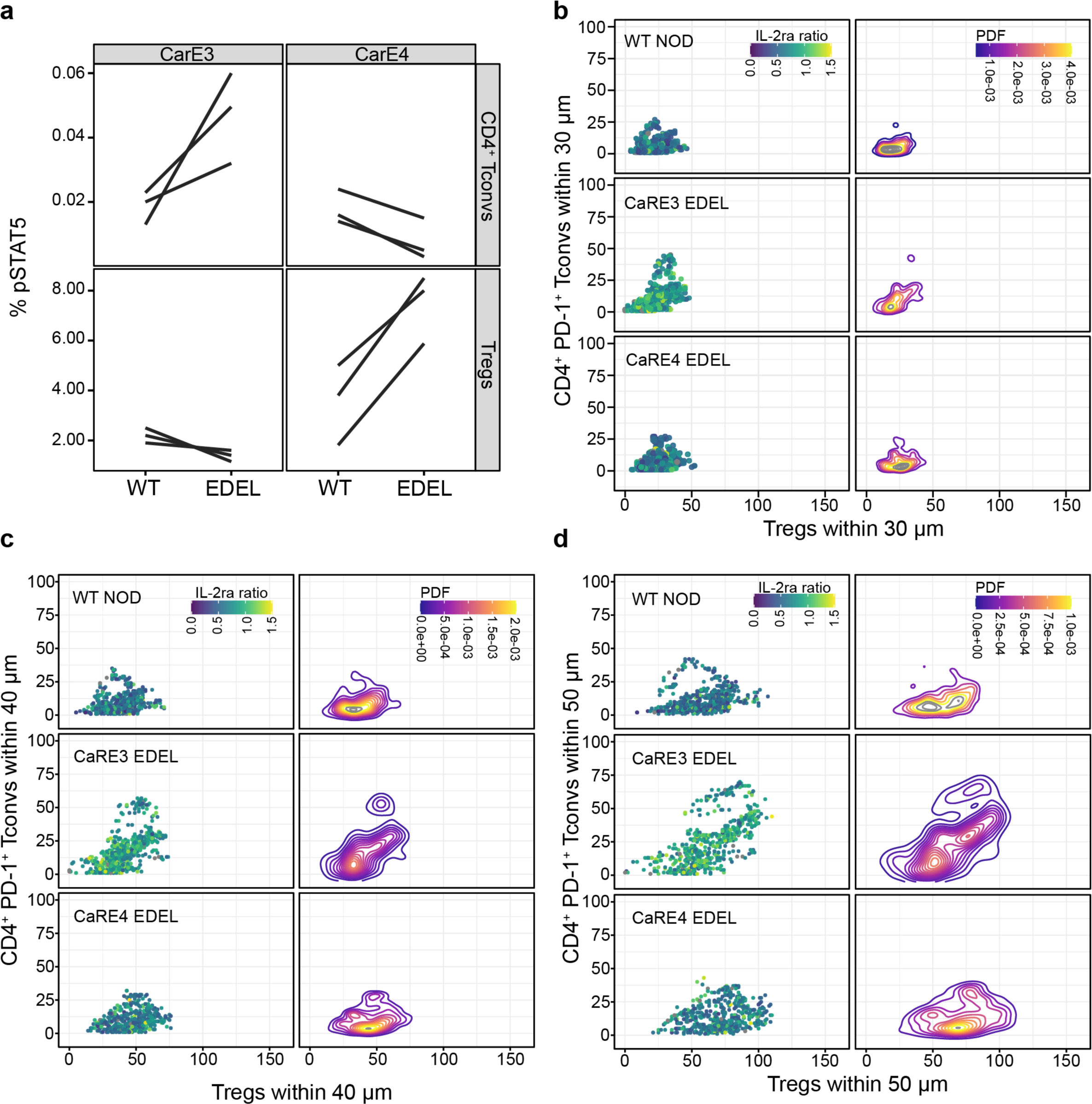

**Supplementary Fig. 6.**
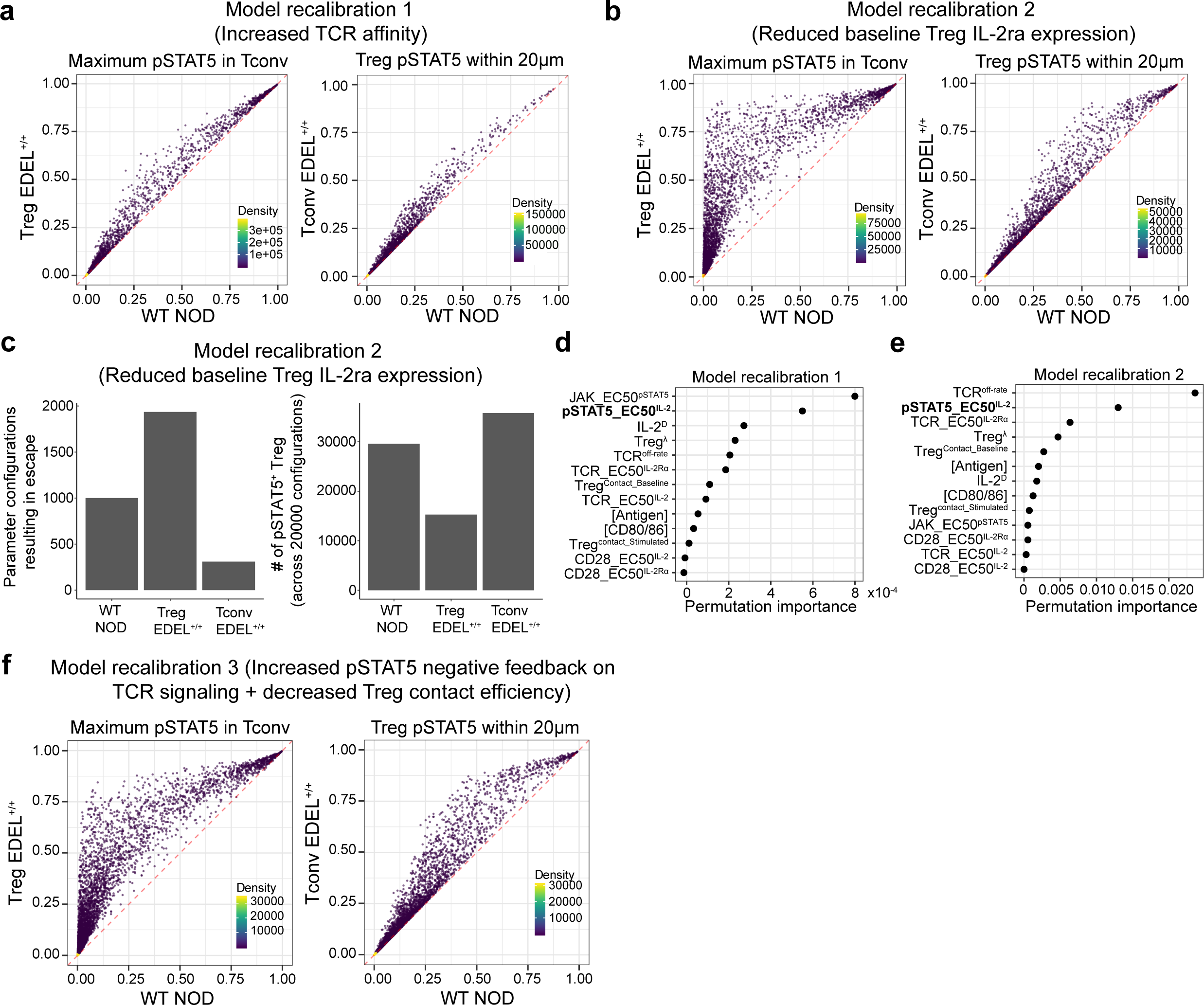

**Supplementary Fig. 7.**
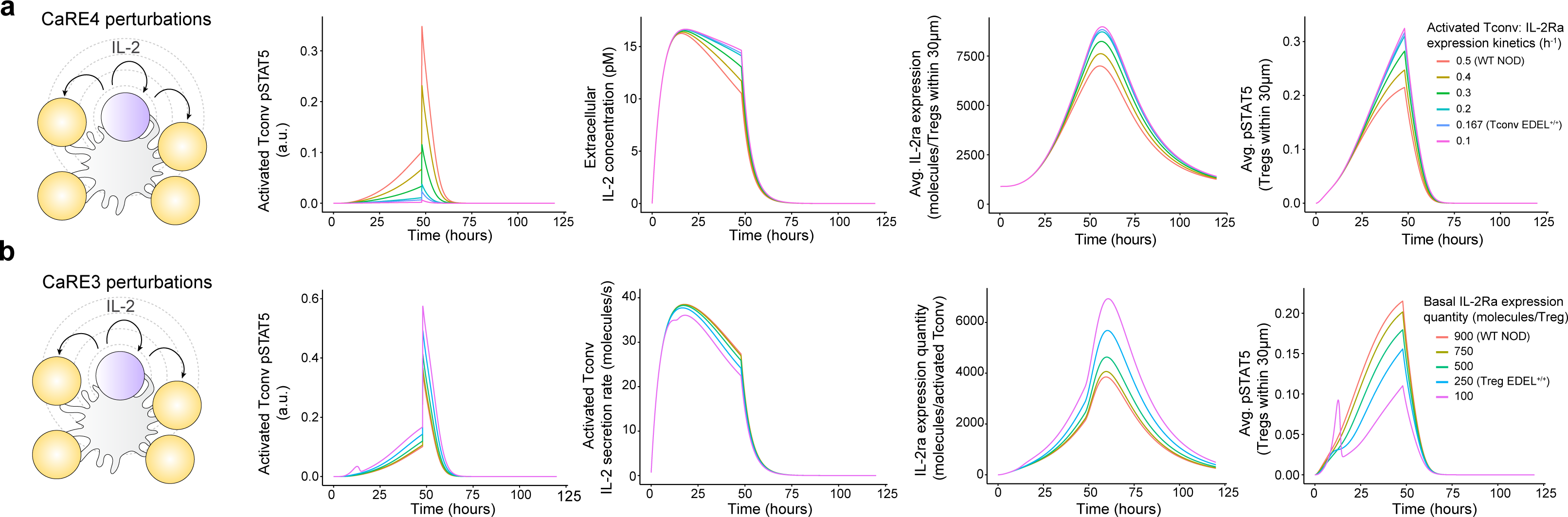

